# Fast ventral stream neural activity enables rapid visual categorization

**DOI:** 10.1101/017897

**Authors:** Maxime Cauchoix, Sébastien M. Crouzet, Denis Fize, Thomas Serre

## Abstract

Primates can recognize objects embedded in complex natural scenes in a glimpse. Rapid categorization paradigms have been extensively used to study our core perceptual abilities when the visual system is forced to operate under strong time constraints. However, the neural underpinning of rapid categorization remains to be understood, and the incredible speed of sight has yet to be reconciled with modern ventral stream cortical theories of shape processing.

Here we recorded multichannel subdural electrocorticogram (ECoG) signals from intermediate areas (V4/PIT) of the ventral stream of the visual cortex while monkeys were actively engaged in detecting the presence or absence of animal targets in natural scenes. Using multivariate pattern analysis (MVPA) techniques, we quantified at millisecond precision task-relevant signals conveyed by ECoG data. Reliable neural decoding was possible shortly after stimulus onset from single trials with a degree of generalization to experimental manipulations closely mimicking monkeys’ accuracy and reaction time.

Together, the present study suggests that rapid ventral stream neural activity induces a selective task-relevant signal subsequently used to drive visual categorization.

**Classification:** Biological sciences / Neuroscience

## INTRODUCTION

The robust and accurate categorization of natural object categories is critical to survival, as it allows an animal to generalize many properties of an object from its category membership [1–5]. Human and non-human primates excel at visual categorization: They can rapidly and reliably categorize objects embedded in complex natural visual scenes in a glimpse [see 6,7 for recent reviews].

It is well known that object recognition is possible for complex natural scenes viewed in rapid visual presentations that do not allow sufficient time for eye movements or shifts of attention [8–10]. The underlying visual representation remains relatively coarse as participants frequently fail to localize targets that are correctly detected in an image stream [11]. Studies using backward-masking [12] and saccadic responses [13,14] have further demonstrated that recognition is possible under severe time constraints. While much is known about the psychological basis of rapid categorization, much less is known about the underlying neural processes and, in particular, the timing of the corresponding perceptual decisions.

Using human scalp electroencephalography (EEG), Thorpe et al. [10] first demonstrated that a category-selective signal can be isolated from frontal electrodes shortly after a stimulus is flashed (within ∼150 ms post stimulus onset). Previous work using intra-cranial recordings has shown that it is possible to decode object category information from the ventral stream of the visual cortex very rapidly (within ∼ 100 ms post stimulus onset) in both humans [15] and monkeys [16–18]. However, this work either used a passive viewing paradigm [15,17,18] or involved a relatively simple basic-level categorization task, such as trees vs. objects [16], and did not establish any link between ventral stream neural activity and (speeded) behavioral responses during rapid categorization. Indeed, previous monkey electrophysiology work has found little co-variation between reaction time and neural latencies in the inferotemporal cortex [19,20,but see also 21].

Here we recorded ECoG activity in monkeys actively engaged in a rapid categorization task with natural scenes. Using multi-variate pattern analysis (MVPA) techniques, we were able to isolate an initial task-relevant neural activity in the ventral stream shortly after stimulus onset. We further assessed how well this early category-selective neural signal mimics behavioral responses in two experimental manipulations: When a backward mask is presented immediately following the stimulus, and when participants are required to generalize beyond familiar exemplars to novel unpredictable stimuli. In both cases, we find that neural decoding closely mimics the monkeys’ behavioral responses.

Overall the present study suggests a link between categorical activity in the ventral stream and behavior, and offers a parsimonious account of rapid visual categorization reconciling the incredible speed of sight with modern ventral stream theories of object recognition [22].

## MATERIAL AND METHODS

Further details about the protocol, the stimuli set, neurophysiological and behavioral data processing and computational approaches models used can be found in the Supporting Information.

### Behavioral testing and data acquisition

Two male rhesus macaques (M1 and M2, both aged 14) were trained to perform a rapid natural scene categorization task by releasing a button and touching the screen when they saw an animal in the stimulus presented (target) or keeping their hand on the button otherwise (distractor). The image set consisted of natural gray-scale photographs (256 □ 256 pixels) equalized for average luminance and global contrast (root mean square over pixel intensities). We report behavioral accuracies as d’ (d’(t)=z[Hits(t)] - z[False Alarms(t)], where z[] is the inverse of the normal distribution function).

Monkeys were implanted with subdural macro-electrodes (Fig S1). Recording was performed using the NeuroScan EEG system and SynAmps amplifier system sampling at 1000 Hz (band pass 0.1-200 Hz). Frontal electrodes were used as reference. The signal was baseline corrected [-50; 30 ms] trial by trial and down-sampled to 512 Hz. All procedures conformed to French and European standards concerning the use of experimental animals. Protocols were approved by the regional ethical committee.

### Data analysis

All neural data analyses were conducted using the Matlab EEGlab toolbox (50) and custom software. We used a procedure similar to Yoshor el al. (23) to map out the visual receptive fields in one animal (M1) and compute ERP visual latencies. MVPA was performed directly on the neural signal using a linear Support Vector Machine (SVM) classifier. MVPA results correspond to the average accuracy computed using a cross-validation procedure (n=300) whereby different training and test sets were selected each time at random. A measure of chance level was obtained by performing the same analysis on permuted labels. The decoding was considered above chance when 95% of the differences between decoding on true or permuted labels paired over the 300 cross-validations fell above zero. In the case of temporal decoding, to correct for multiple comparisons, a bin was considered significant when followed by at least five consecutive significant bins, (p <0.01) as done in Liu et al. (Liu et al., 2009).

## RESULTS

### Successful learning of the categorization task

Two monkeys (M1 and M2) were trained to perform a rapid categorization task in which they had to report the presence or absence of an animal in briefly presented natural scenes (Fig. 1). Despite the large variability of the natural stimuli used (Fig. 1a), both monkeys were able to learn to categorize images with a high degree of accuracy (measured as d’; M1: 3.15; M2: 2.96) and very short median reaction times (RTs; M1: 305 ms; M2: 289 ms).

**Figure 1.**
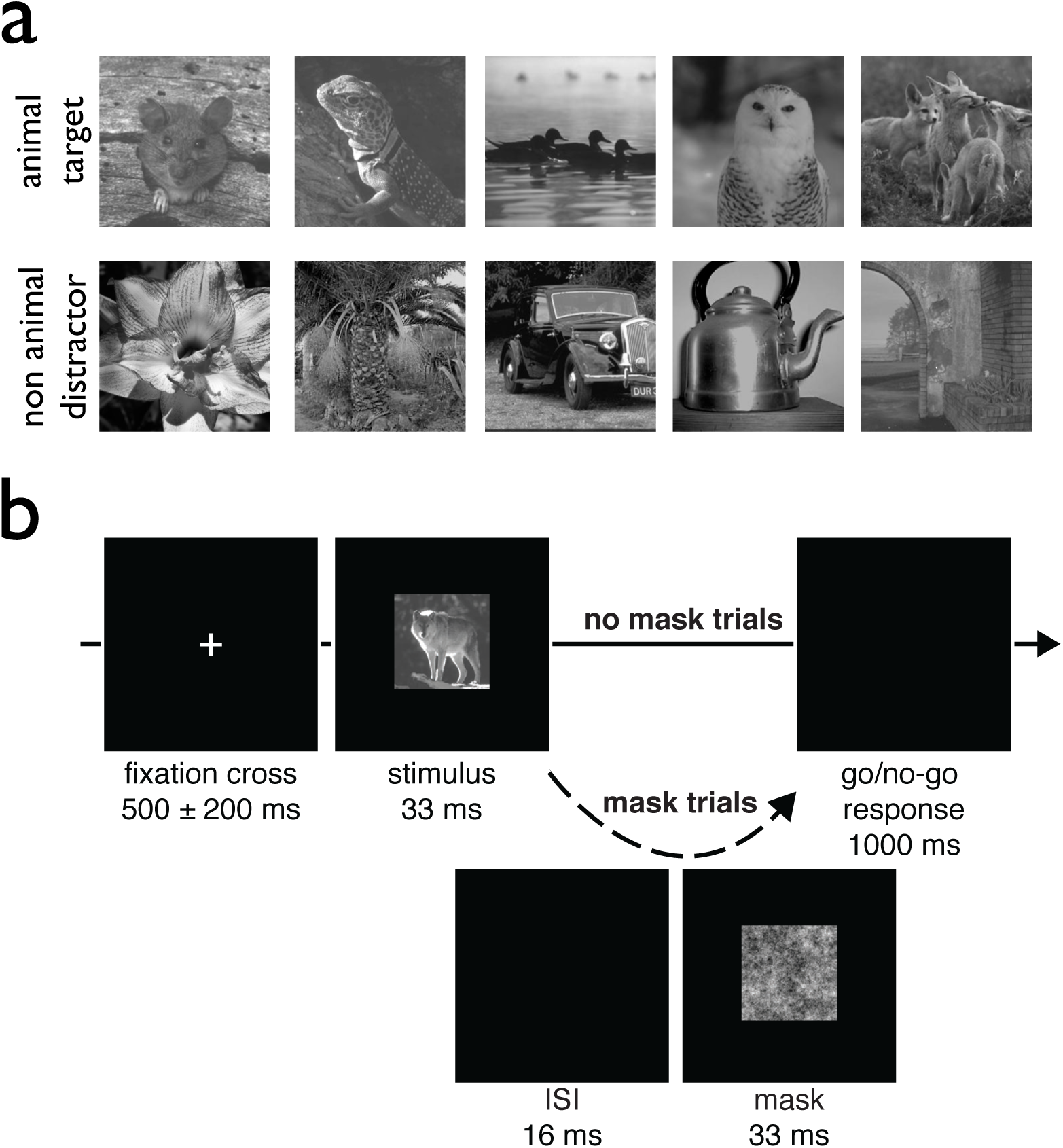
Stimulus set and visual categorization task. **a)**The stimulus set consists of natural gray-scale images with both animal targets (n=340) and non-animal distractors (n=340). **b)** Following the presentation of a fixation cross, an image is flashed for 33 ms. The monkeys indicate whether a target is present by releasing a button within 1 s following image presentation. On half the trials, a backward mask (1/f pink noise) is displayed for an additional 33 ms following a 16-ms blank screen (50 ms SOA).

### Event-Related Potentials (ERPs) and visual receptive fields

While the two animals were engaged in the rapid categorization task, we recorded subdural electrocorticogram (ECoG) signals from multiple electrodes implanted over intermediate areas (V4/PIT) of the visual cortex (Fig. 2, FigS1). We estimated the visual latency of individual electrodes using the method proposed by Yoshor et al. [23]. Visual latencies ranged between ∼40–70 ms post stimulus onset for M1 (median: 53 ms) and from ∼35–80 ms for M2 (median: 59 ms), with a trend for V4 electrodes to exhibit shorter latencies than PIT electrodes (Fig S2).

**Figure 2.**
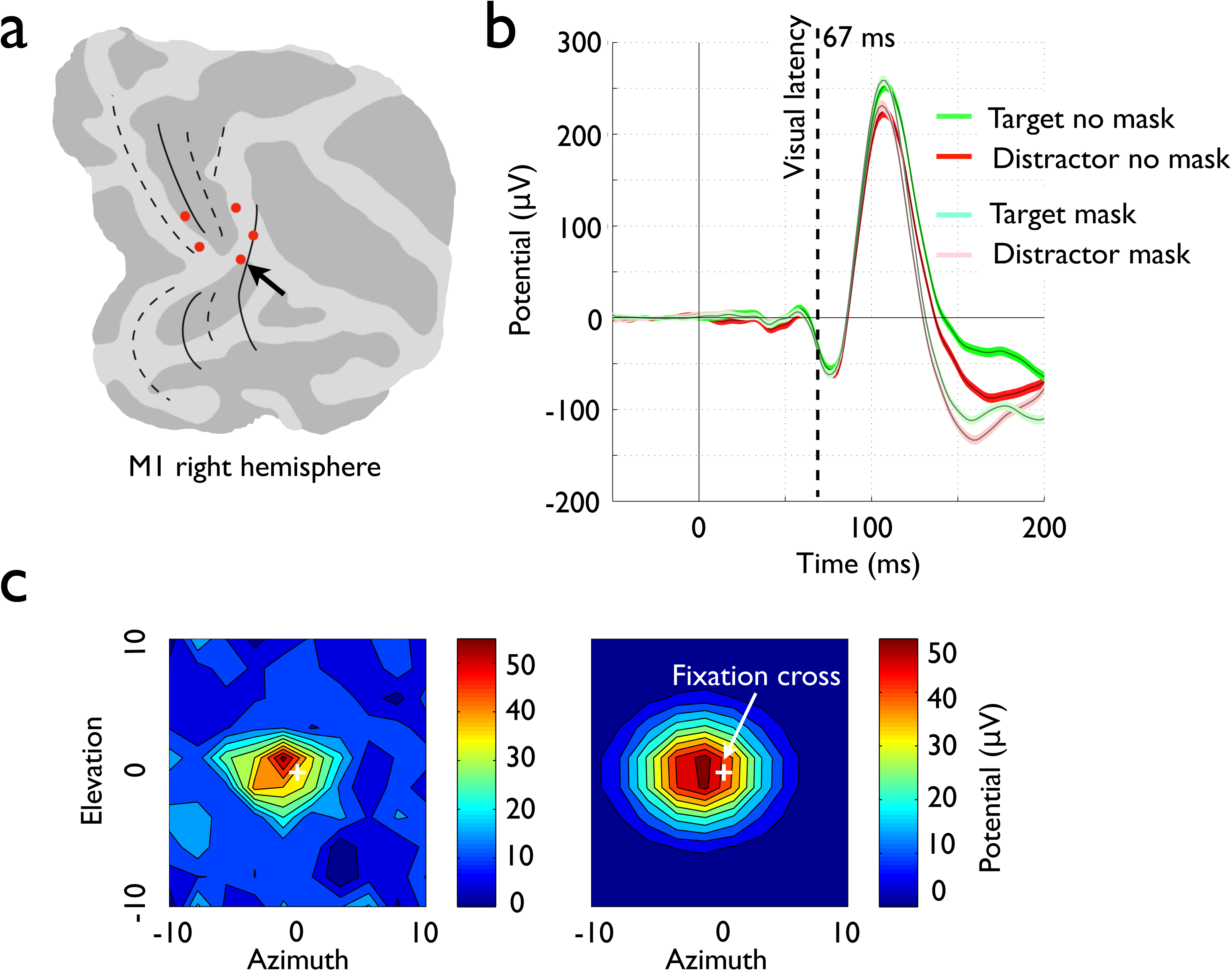
Sample electrode locations, Event Related Potential and visual receptive field. **a)** Electrode tips (red dots) are shown on (FreeSurfer) flat maps for monkey M1 (right hemisphere). **b)** The corresponding ERP for the electrode marked with an arrow in panel a. Shown is the average potential evoked by animal targets (green) vs. non-animal distractors (red) for both the mask (darker shade) vs. no-mask (lighter shade) conditions. Thin black lines indicate sample averages with corresponding CIs (95%) obtained via bootstrapping shown with transparency. **c)** Visual receptive field for the same electrode before (left) and after (right) two-dimensional Gaussian fitting.

To verify that the recorded neural responses were visual in nature, we mapped out receptive fields (RFs) coarsely in one of the two animals (M1) by flashing small white squares during the fixation-cross (pre-trial) intervals (see Methods). RF sizes ranged between 3.4°–8.9° with an average of 7.0° and were of similar size for V4 and PIT electrodes (Fig. 2c, Fig S3).

We further estimated the earliest significant differential activity for animal vs. non-animal stimuli for each individual electrode using a point-by-point analysis as done in [10,15]: Latency here is defined as the first time point where five consecutive points (10 ms bins) yields p < 0.01 in a t-test. For one of the two monkeys (M1), all electrodes exhibited a significant differential activity for animal vs. non-animal with a latency under 200 ms. For the other monkey (M2), only about half of the electrodes (8 out of 13) exhibited significance. The earliest significant animal/non-animal differential activity found using this method occurred at 83 ms in M1 on one V4 electrode and at 89 ms in M2 for a PIT electrode (Fig S2).

### Fast single-trial decoding of superordinate category

We further assessed how well category information can be decoded from across all electrodes using a Multi-Variate Pattern Analysis (MVPA). We trained and tested a linear Support Vector Machine (SVM) classifier directly on pooled electrode potentials for every time point independently. This type of MVPA typically provides higher statistical power over univariate methods by pooling information across all electrodes, enabling reliable estimates of latencies from single trials [24]. Here we found that, indeed, reliable decoding of superordinate category information (animal vs. non animal) was possible from single trials under 100 ms post stimulus onset (M1: 92 ms, M2: 96 ms; Fig. 3a) and that electrodes from both V4 and PIT contributed to the decoding accuracy (Fig. S4).

**Figure 3.**
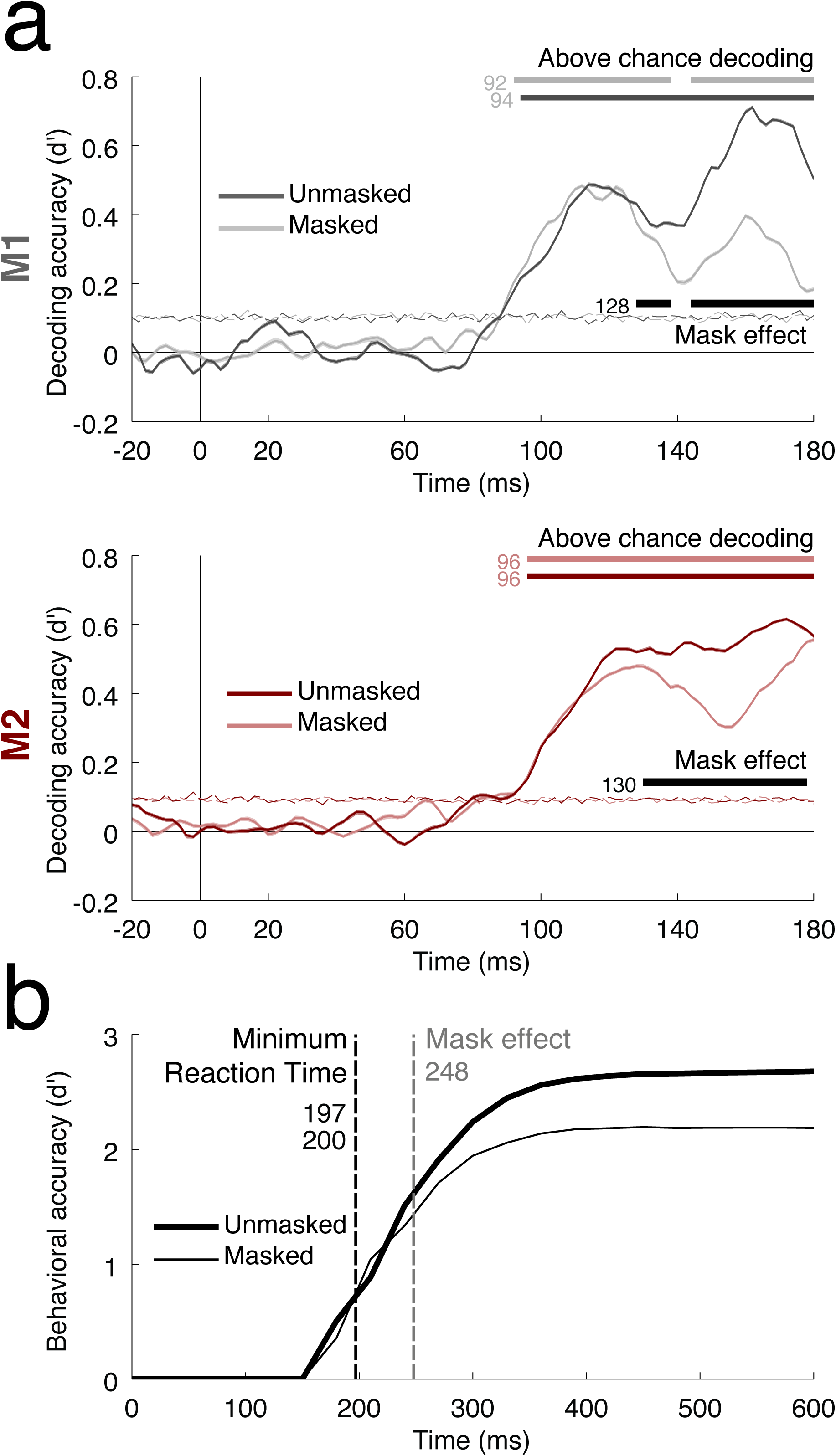
Decoding from ventral stream neural activity and backward mask effect. **a)** Readout accuracy for monkey M1 (gray) and M2 (red) during the mask (lighter shade) vs. no-mask conditions (darker shade). Center lines correspond to the average decoding accuracy estimated across multiple cross-validations of the data. Corresponding CIs (95%) obtained via bootstrapping are shown with transparency (most confidence intervals are actually too small to be visible). Horizontal lines indicate decoding latencies for the mask and no-mask conditions (upper colored bars) as well as the earliest significant effect associated with the presentation of the mask on decoding accuracy (lower black bar on each plot). Horizontal dotted lines indicate the upper limit of the 95% CI around chance level obtained with a permutation procedure. **b)** Cumulative d’ curves plotted as a function of response times for mask (thin) vs. no-mask conditions (thick). Black dotted vertical line indicates minimum reaction times for mask (thin) vs. no-mask conditions (thick). Gray vertical line indicates earliest mask effect.

Next, we assessed how well this early category-selective neural signal mimics behavioral responses in two experimental manipulations: (1) When a backward mask is presented immediately following the stimulus, and (2) when the animals are required to generalize beyond familiar exemplars to novel unpredictable stimuli.

### Backward masking alters behavior and ECoG decoding similarly

On half of the trials, a backward mask (1/f pink noise) was presented following stimulus presentation with a stimulus onset asynchrony (SOA) of 50 ms. This type of mask puts very severe time constraints on the visual system: It is assumed to interrupt visual processing by disrupting visual persistence and hindering recurrent signals [25,26], thus encouraging fast, i.e. mostly feedforward, behavioral responses [27]. Consistent with this hypothesis, previous human psychophysics studies have established that backward masking with an SOA ∼50 ms (as used here) or longer has very little effect on the fastest human behavioral responses [27] and early scalp electro-encephalography (target vs. distractor) differential activity [12].

Here, the overall behavioral accuracy (d’) of the two animals (computed on both familiar and novel images) was significantly reduced on masked trials (M1: 2.12 vs. 2.83 for mask vs. no-mask presentations, χ2(1)=99, p<0.001; M2: 2.14 vs. 2.55 for mask vs. no-mask presentations, χ2(1)=40, p<0.001) but far exceeded chance level (M1: χ2(1)=2059, p<0.001; M2: χ2(1)=2326, p<0.001).

Cumulative d’ curves plotted as a function of response times are shown in Fig. 3b. The cumulative number of hit and false alarm responses was used to calculate an accuracy measure at each time point t. Such analyzes of the time course of reaction times aim to provide a behavioral characterization of the underlying processing dynamics. The shortest RT was unaffected by the presentation of the mask (min RT = 197/200 ms with/without mask). A significant difference between masked and unmasked trials only appeared for trials with slower responses (with a delay roughly equal to the SOA ∼50 ms; Fig. 3b).

An ECoG decoding analysis performed separately on trials with/without masking revealed very similar estimates of decoding latencies (M1: 94 ms, M2: 96 ms). Interestingly, the first significant difference between the two conditions was found during a later time window (starting at 128 ms post stimulus onset for M1 and 130 ms for M2; Fig. 3a). This result matches well with the behavioral results reported above, and suggests that the mask appears to leave unaffected the first 40-50 ms of visual processing.

### Fast ventral stream neural activity enables generalization to novel exemplars

We then evaluated behavioral responses and neural decoding accuracy separately for a novel and familiar set of images. On the very first presentation of the novel stimuli (80 images x 5 days), the accuracy of the monkeys (d’) remained well above chance (M1: 1.62, χ2(1)=96, p<0.001; M2: 1.99, χ2(1)=123, p<0.001). Similarly, global decoding from early ventral stream neural activity (90–140 ms time window post-stimulus onset) generalized well above chance from the familiar (M1: 0.68, p<0.01; M2: 0.79, p<0.01) to the novel set (M1: 0.34, p<0.05; M2: 0.48, p<0.05).

These results suggest that the two monkeys did form a high-level concept of the animal category beyond rote generalization. This is all the more remarkable as the stimulus dataset used includes multiple animal species (mammals, reptiles, fishes, birds, etc.) and multiple factors that can affect the appearance of the target object category such as changes in position, scale, view-point or the type of background scenery. Rapid ventral stream neural activity may thus play a key role in supporting primates’ generalization ability.

### Fast single-trial decoding is predictive of behavioral responses

To further link this early ventral stream neural activity to behavioral accuracy, we computed an “animalness” accuracy score for individual images based on either monkeys’ behavioral responses or neural decoding (Fig S5). Such a score provides a compact characterization of the visual strategy employed by a visual system and permits the comparison between neural and behavioral data via direct correlation. The correlation between animalness scores computed from neural decoding (90–140 ms time window) and behavioral responses was significant (M1: r^2^=0.45, p<0.001; M2: r^2^=0.27, p<0.001), even after correcting for classification accuracy using partial correlation (see Methods; M1: r^2^*=0.29, p<0.001; M2: r^2^*=0.15, p<0.001).

We computed a similar score for a representative feedforward computational model of the ventral stream of the visual cortex (HMAX) previously shown to match human performance on the same rapid animal categorization task [28]. A significant correlation was found between the HMAX model and monkey behavior (M1: r^2^=0.41, r^2^*=0.16, p<0.001; M2: r^2^=0.41, r^2^*=0.13, p<0.001) as well as neural activity (M1: r^2^=0.28, r^2^*=0.14, p<0.001; M2: r^2^=0.20, r^2^*=0.09, p<0.001). In contrast, no significant correlation was observed between neural activity and a low-level visual representation based on pixel intensities (M1: r^2^□0, r^2^* □0, p=0.8; M2: r^2^=0.02, r^2^*=0.02, p=0.06).

To test for a more direct link between this early neural activity and behavioral response times, a hallmark of decision making processes [29,30], we organized trials into quartiles based on the overall distribution of reaction times (Fig. 4a). Global decoding (90-140 ms post-stimulus onset time window) revealed a monotonic (near-linear) relationship between decoding accuracy and mean reaction times for each quartile (Fig. 4a): Trials corresponding to faster quartiles were decoded with higher accuracy (M1: F(3,1196)=1477, p<0.001; M2: F(3,1196)=820, p<0.001). Temporal decoding conducted on individual quartiles separately (Fig. 4b) further suggested that decoding latencies followed response times very closely. Trials associated with faster quartiles were decoded faster and with higher accuracy.

**Figure 4.**
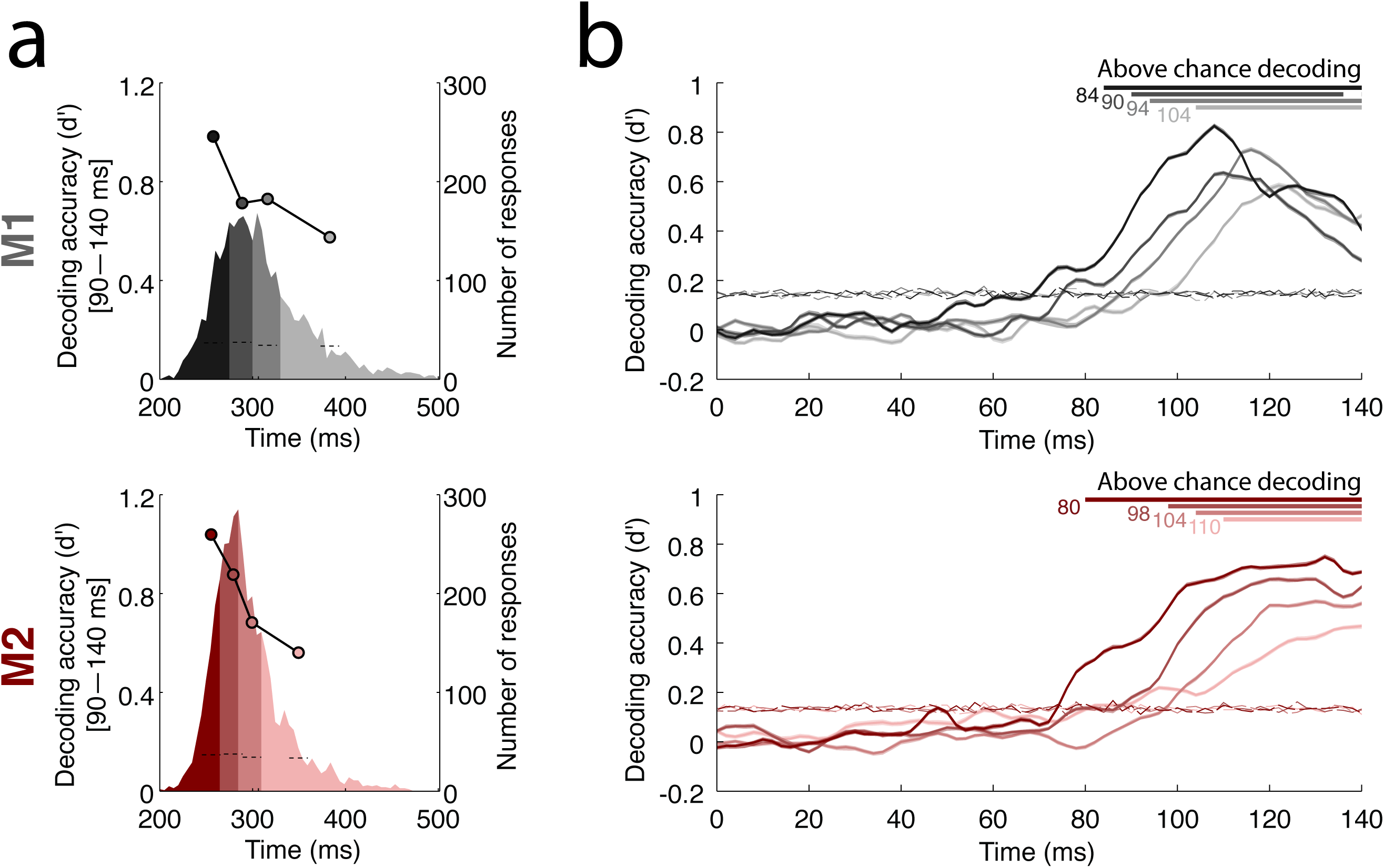
Linking rapid ventral stream neural activity with response times.. **a)** Trial binning according to reaction times and corresponding neural decoding (overlaid on the distributions) shown as circles (90-140 ms time window; 95% CI shown as error bars) for monkey M1 (gray) and M2 (red). Horizontal dotted lines indicate upper limit of the 95% CI around chance level obtained with a permutation procedure. b) Decoding conducted on individual quartiles. Center lines correspond to the average decoding accuracy estimated over multiple cross-validations of the data. Corresponding CIs (95%) obtained via bootstrapping are shown with transparency (most confidence intervals are actually too small to be visible). Significant deviation from chance level is shown at the top of the graph with horizontal bars along with the corresponding earliest latencies. Horizontal dotted curves indicate upper limit of the 95% CI around chance level obtained with a permutation procedure.

Overall, these results demonstrate that fast neural activity in the ventral stream is linked to both behavioral accuracy and reaction time.

### Fine visual information can be decoded independently of monkey behavior

To further quantify the visual information encoded in this neural activity beyond task-related category information, we considered four (basic) categories of images from a subset of the target stimuli presented (people, macaques, chimpanzees, otter faces; Fig. S6). We trained and tested a classifier to discriminate between these four subcategories of animal images using a one-versus-all decoding procedure. Reliable decoding of these subcategories above chance level suggests that task-related information does not completely override visual information. These results further suggest that, in principle, fast ventral stream neural activity could subserve multiple visual recognition tasks, consistent with earlier predictions from computational models [31,32].

### Human and non-human primates share a similar visual strategy

Here, we compare the accuracy (d’) of the two monkeys to that of human participants [28] using a very similar paradigm with the same novel set of stimuli. Despite the presence of the mask, monkeys reached an accuracy level (M1: 1.35; M2: 1.92) comparable to that of human participants (n=22; H: mean 1.96, SD=0.50) for the same novel set of images.

To further assess the similarity of the visual strategies used by monkeys and human participants, we computed animalness scores for both from behavioral responses (Fig. S7). For human observers, this score was computed as the fraction of human observers that classified a specific image as an animal irrespective of the actual category label. A score of 1.0/0.0 means that all participants classified the image as animal/non-animal. Any value in between reflects some variability across subjects. We computed a similar index for monkeys by pooling responses between the two animals and over multiple trials to obtain a reliable estimate.

We found a significant correlation between human observers and monkeys (Spearman correlation: r^2^=0.73, p<0.001) even after factoring out accuracy using a partial correlation measure (r^2^*=0.33, p<0.001). This interspecies correlation was as strong as the correlation between the two monkeys (r^2^=0.67, p<0.001; partial correlation: r^2^*=0.32, p<0.001). Overall this suggests that monkey and human participants do indeed follow a very similar visual strategy in our task.

## DISCUSSION

The present study investigated the neural underpinning of rapid natural scene categorization in non-human primates. Despite the inherent complexity of natural scenes, we found that superordinate object category can be read out very rapidly (within 100ms post stimuli onset) from intermediate areas (V4/PIT) of the ventral stream of the visual cortex. One limitation of ECoG recordings is the lack of a precise spatial localization of the underlying neural source. We remain confident, however, that the recorded neural signals were not contaminated by motor preparatory responses because our analysis time window did not overlap with behavioral responses. Furthermore, in addition to task-relevant category signals, we were also able to decode task-irrelevant visual information and to map out RFs for individual electrodes.

Using backward masking, we found a striking degree of similarity between ventral stream neural activity and behavioral responses: both remained unaffected during an initial visual processing stage with the mask only impacting later processing. These results are consistent with earlier electrophysiological masking studies [26,33,34] and masking theories [35,36] positing that visual processing of the stimulus and mask are kept separate during an initial short time period. These results are also consistent with theories postulating two distinct modes of visual processing, i.e., an early (possibly feedforward) processing unaltered by the presentation of a backward mask, which only interferes with later (feedback) processing [25,27,37]. At the same time, given our relatively short estimate of the timing of the mask effect (∼130 ms) compared to estimates of V4 attentional modulation reported in the literature (∼160–170 ms, see [38,39]), the presentation of the mask in our study is likely to already interfere with feedforward processing. In addition, the observed mask timing does not exclude a possible fast recurrent modulation during the initial processing time window [40]. A more direct test for teasing apart alternative theories of visual masking would, however, require to vary the SOA more systematically to demonstrate co-variations between SOA, neural decoding and behavior, or the use of an inactivation protocol as done in Hupé et al. [41].

Most previous attempts to link visual processes to behavior have focused on the processing of motion information in the dorsal stream of the visual cortex [42]. A previous study based on single unit activity in IT did not find any co-variation between neural latencies in IT and reaction time [20]. The present study using ECoG electrode, however, was able to identify co-variations between ventral stream neural activity and both perceptual decisions and their timing. This is consistent with a more recent single-cell study which demonstrated a correspondence between neural activity in IT and the speed of recognition [21] using isolated objects in a visual search display. Furthermore, the estimated latency of category information overlaps with the optimal timing for micro-stimulations in IT to affect perceptual decisions [43].

Our analysis further suggests that relatively modest shifts in the latency of task-related information encoded in the ventral stream (4–18ms) yields larger shifts in the corresponding distribution of behavioral responses (10–60ms). One possible interpretation based on computational models of decision making such as the Drift Diffusion Model (DDM; [44]) is that ventral stream neural activity reflects the rate of accumulation of information, known as the drift rate, which is determined by the quality of the information extracted from the visual stimulus. The role of the ventral stream would thus be to convey the amount of evidence in the stimulus toward the target or the distractor decision category [44].

The impressive speed of visual processing observed during rapid categorization tasks has led some researchers to argue for subcortical routes bypassing altogether the visual cortex, e.g., via the thalamus through the amygdala [45]. Direct projections between these two structures were observed on rodent electrophysiology during fear conditioning, but evidence for this “low-road” subcortical pathway for rapid vision is a matter of debate [see 46 for a recent review]. The existence of animal category selective responses has been demonstrated in the human amygdala [47]. The observed latencies are relatively late (>300 ms) compared to those found in the present study (<100 ms). Interestingly a recent monkey electrophysiology study has shown selectivity for threatening stimuli (snakes) in the pulvinar within ∼50 ms post-stimulus onset [48]. The present study supports more directly the “high-road” (cortical) hypothesis. At the same time, the existence of direct projections between the pulvinar and intermediate areas of the ventral stream of the visual cortex leaves, however, open the possibility that the visual selectivity for animal vs. non-animal found in the present study originates in subcortical areas with the ventral stream simply relaying the information to downstream areas [49].

Evidence suggesting that monkeys are capable of learning a high-level visual concept of abstract natural categories has so far been limited to behavioral studies [5,50]. Previous monkey electrophysiology studies on categorization (e.g., cats vs. dogs [51] or faces vs. fishes [52]) did not test for generalization to novel (unfamiliar) stimuli and are thus compatible with rote learning of individual exemplars [but see also 16,53]. The present study establishes that ventral stream neural activity, in principle, could be supporting primates’ generalization ability during high-level processing of natural scenes. In addition, we also found that it was possible to decode finer level (basic) category information, which was unrelated to the task (from target stimuli only). This suggests that, while salient, task-related category information does not completely override the visual information contained in these relatively early visual processes.

We further demonstrated that monkey and human participants exhibit similar patterns of correct and incorrect responses on the same set of images suggesting that they engage similar visual representations. These behavioral results adds to a growing body of evidence [5] suggesting that the neural mechanisms supporting rapid object categorization may be conserved between humans and macaque monkeys.

In addition, the present study found a high degree of correlation between neural and behavioral data as well as a computational model of the ventral stream of the visual cortex. While we controlled for the most obvious low-level visual differences between the target and distractor set (e.g., distance to the camera, pixel intensities), we cannot rule out the possibility that relatively low-level features (including spatial frequencies) may be driving these correlations.

In sum, the present study suggests that rapid ventral stream neural activity induces a selective task-relevant signal subsequently used to drive visual categorization.

## ACKNOWLEDGMENT

We would like to thank several of our colleagues for their valuable inputs on this manuscript: N. Bichot, D. Brooks, M. Franck, M. Peelen, T. Poggio, D. Sheinberg, I. Sofer and R. VanRullen. The monkey electrophysiology was supported by the FRM and DGA. The analysis of the data and comparison to models was supported by NSF early career award (IIS-1252951) and DARPA grant (N10AP20013) to T.S. Additional support to T.S. was provided by ONR grant (N000141110743), the Center for Computation and Visualization at Brown University, and the Robert J. and Nancy D. Carney Fund for Scientific Innovation.

